# Submaximal running energetics are maintained despite local muscle fatigue

**DOI:** 10.64898/2026.01.16.699896

**Authors:** Key Nahan, Harrison Finn, John Kerr, Martin Héroux, Kirsty A. McDonald

**Author notes:** Correspondence: *Key Nahan (*).

## Abstract

When running, metabolic cost increases as muscles are simultaneously fatigued. However, the contribution of an individual muscle group to fatigue-related increase in metabolic costs remains unclear. We investigated the metabolic consequence of running with local plantar flexor or knee extensor fatigue and associated neuromuscular control strategies. *Recreational* and *experienced* male runners (N=20) completed two sessions (one per muscle group), with each including two 10 min running bouts: without and with local fatigue (∼20% reduction in peak joint torque). Net metabolic power and muscle activity (initial and final minutes) were determined. Metabolic power was unaffected by plantar flexor (p=0.367) or knee extensor (p=0.607) fatigue in both cohorts. Plantar flexor fatigue recovered during the *fatigued run* (p=0.033), while knee extensor fatigue only recovered for the *recreational* cohort (p=0.009; *experienced*: p=0.826). With plantar flexor fatigue, plantar flexor muscle activity was unchanged between runs (p≥0.312), however initial soleus activity was greater in the *unfatigued* than *fatigued run* for *experienced* runners (p=0.022), and initial medial gastrocnemius activity was greater in the *unfatigued* than *fatigued run* for the combined cohort (p=0.009). With knee extensor fatigue, knee extensor muscle activity was mostly lower in the *unfatigued* than *fatigued run* (p≤0.009), except for final vastus lateralis activity, which was unchanged between runs (p=0.061). Therefore, muscle groups respond with different activation strategies when fatigued. Running with plantar flexor or knee extensor fatigue, at levels like those induced by prolonged running (10-42 km), does not increase metabolic power and thus, submaximal running energetics may be maintained despite local muscle fatigue.

**NEW & NOTEWORTHY:** While muscle fatigue is suggested to increase the metabolic cost of running, the individual contributions of key lower limb muscle groups have not been explored. We examined responses after fatigue of only the plantar flexors or the knee extensors. Results indicate that local fatigue did not affect the metabolic power of male runners for either fatigued muscle group. These findings enhance our understanding of running performance and the interaction between fundamental criteria dictating human locomotion.

## INTRODUCTION

The metabolic cost of running, as represented by metabolic power or cost of transport, is a comprehensive measure of submaximal energy consumption and a primary determinant of running performance (1–3). Neuromuscular fatigue (i.e., reduced maximal force production of a given muscle (4)), a secondary determinant, may increase metabolic cost (5, 6), reduce running speed (5, 7, 8) and subsequently influence running performance. For example, a fatiguing squat protocol increases the metabolic cost of running (9), while a fatiguing drop jump protocol increases metabolic cost (10) and time to exhaustion (11) in cycling. Moreover, lower fatigability of the knee extensors has been associated with preserved running economy when measured before and after a fatiguing run (12).

Physiological and biomechanical mechanisms underpin the inflated metabolic cost of running with neuromuscular fatigue. Physiologically, an increased recruitment of larger, type II muscle fibers during fatiguing contractions (13, 14) may contribute to the metabolic penalty as they require more energy to contract (15–18). A change in the substrate used when muscles are fatigued may also contribute, as reliance on glycogen decreases and free fatty acid increases (19, 20). During aerobic exercise, adenosine triphosphate (ATP) produced from fat is approximately 7% less efficient than that from carbohydrates (21); thus, the metabolic cost of running may be affected by changing fuel sources. Similarly, neuromuscular fatigue is associated with increased metabolite (i.e., hydrogen ion, inorganic phosphate) concentrations; this may decrease intracellular pH and result in a reduced rate of ATP production and increased metabolic cost (22, 23). Biomechanically, changes in tissue mechanics as a result of neuromuscular fatigue (e.g., increased tendon compliance (24, 25)) may increase the metabolic cost of running (26, 27). Reduced stride frequency (28–30) and gait alterations to offload the fatigued muscle (31) may also contribute to a less metabolically optimal gait strategy and impaired running performance (32, 33).

Running-induced neuromuscular fatigue is difficult to quantify. A run induces fatigue across many muscles simultaneously (i.e., global fatigue) and to varying extents. Functional strength tests, such as drop or countermovement jump tests, do not measure neuromuscular fatigue in a single muscle group (34, 35). Alternatively, muscle activity, measured via electromyography (EMG), increases with fatigue (36) due to increased motor unit firing rate, additional recruitment of type II muscle fibers and/or synchronization of motor unit firing (36). Although EMG provides insight into neuromuscular responses to fatigue, it remains an inaccurate measure of fatigue (37). A maximum voluntary contraction (MVC) is typically used to measure neuromuscular fatigue (38). However, because partial fatigue recovery occurs almost immediately after exercise is stopped, with substantial recovery occurring in the first 1-2 min (39, 40), only one muscle group is often able to be measured after running. Therefore, assessing fatigue in a single muscle group (i.e., local fatigue) via an MVC provides the most accurate measurement of neuromuscular fatigue, effectively circumventing measurement barriers.

The plantar flexors and knee extensors are the main contributors to center of mass acceleration (i.e., positive work) and deceleration (i.e., negative work) in running (41). Considering that muscle work explains ∼76% of the metabolic cost of running (42), accumulation of fatigue in these muscle groups likely inflicts a metabolic penalty. After a prolonged run (10-42 km), peak torque decreases by 9-29% in the plantar flexors (6, 43–47) and 12-30% in the knee extensors (6–8, 35, 48–51). However, it is not possible to determine the metabolic cost of running with such levels of fatigue because i) running-induced fatigue protocols cause neuromuscular fatigue in several lower limb muscle groups and ii) metabolic cost increases following a prolonged run can result from several factors in addition to neuromuscular fatigue (e.g., heat dissipation (52), pulmonary ventilation (53), and heart rate (54)). A local neuromuscular fatigue protocol that consists of a repeated single joint movement, e.g., calf raise or knee extension, may be better suited to explore the metabolic cost of neuromuscular fatigue in specific lower limb muscle groups. Indeed, this type of protocol has been used to explore the effect of local neuromuscular fatigue on running biomechanics (55, 56). While biomechanical changes were observed, it remains unclear whether this fatigue also incurs a metabolic penalty. To address this knowledge gap, we aimed to quantify the effect of local i) plantar flexor and ii) knee extensor fatigue on metabolic cost and fatigued-group muscle activity in running. First, we hypothesized that the metabolic cost of running would increase with local fatigue of either muscle groups. Second, because muscle activity of the plantar flexors and knee extensors increases with running-induced fatigue (5, 57), we hypothesized that local fatigue would also increase activity in fatigued muscles when running.

## MATERIALS AND METHODS

### Participants

All study procedures were approved by the University of New South Wales Human Research Ethics Committee (HC220753). Healthy male participants (N=20; age: 27.2 ± 5.7 years; body mass: 74.6 ± 9.0 kg; height: 1.80 ± 0.07 m; mean ± SD) gave their written informed consent to participate in the study after clear explanations of study procedures were provided. Females have less fatiguability than males (58), and effort and fatiguability are influenced by the specific phase of the menstrual cycle (59), with these effects often being unique to the individual (60). Therefore, we recruited only male participants to limit confounding factors.

Participants were grouped as *recreational* (N=11; age: 26.9 ± 4.9 years; body mass: 78.1 ± 8.8 kg; height: 1.81 ± 0.08 m) or *experienced* (N=9; age: 27.6 ± 6.7 years; body mass: 70.4 ± 7.5 kg; height: 1.79 ± 0.05 m) according to their 10 km running performance and training habits. *Recreational* participants were required to have a 10 km run time between 50-60 min and were also required to have run 10-20 km/week, 2-4 times per week over the past six months. *Experienced* participants were required to have at least three years of running experience, a 10 km run time of 43 min or less and were also required to have run at least 40 km/week, at least three times per week, over the past six months. All participants were free of respiratory, cardiovascular, neurological, or other conditions that could affect gait. Additional criteria required participants not to have experienced any lower limb injuries over the past six months and to have no history of rhabdomyolysis. Details for individual participants are listed in Supplementary Table 1.

### Experimental preparation

The study protocol was part of a two-day analysis of running performance. Plantar flexor and knee extensor fatigue were assessed on separate non-consecutive days, randomly ordered, with an average between-session duration of 23 ± 16 days (mean ± SD; range 6-62 days). On the day before both data collection sessions, participants were asked to refrain from their weekly long run and high-intensity exercise. On the day of data collection, participants were asked to refrain from caffeine and exercise, and to fast for the two hours immediately preceding their laboratory visit (water was permitted). Upon arrival, they completed a rating of fatigue scale to reflect their overall fatigue (61). Next, anthropometric measurements were taken (e.g., body mass, height) and limb dominance was determined by asking the participant “which foot would you kick a soccer ball with?” (62). Surface EMG (Trigno Avanti, Delsys, Natick, MA, USA) sites were prepared for seven muscles on the dominant limb according to the SENIEM guidelines (63): gluteus maximus, biceps femoris, rectus femoris (RF), vastus lateralis (VL), medial gastrocnemius (MG), soleus (SOL), and tibialis anterior. Force-sensing resistor pads (Trigno 4 Contact, Delsys, Natick, MA, USA) were placed under the insole of standardized ASICS GEL-CUMULUS 24^TM^ footwear, in line with the mid-calcaneus and first metatarsophalangeal joint (31). The ASICS GEL-CUMULUS 24^TM^ has a 8 mm drop and 16/24 mm (forefoot/rearfoot) midsole stack of ethylene vinyl acetate (64). Foot strike was not classified.

### Experimental protocol

Participants completed a 5 min seated resting metabolic rate trial where breath-by-breath 𝑉͘_𝑂_2__(L s^-1^) and 𝑉͘_𝐶𝑂2_ (L s^-1^) were collected using a portable metabolic system (K5, COSMED, Rome, Italy). They then performed a 5 min warm-up run on a treadmill (TMX428CP, Trackmaster, Full Vision, Inc, Newton, KS, USA) at 8 km h^-1^ [2.22 m s^-1^], 0° incline. Next, participants completed four 3-5 s isometric MVCs on an isokinetic dynamometer (10 kHz, HUMAC NORM, CSMi, Stoughton, MA, USA) for the muscle group of interest (either plantar flexor or knee extensor, depending on the session) with a 1 min rest between contractions (43, 50). Verbal encouragement was given during the contractions (65). Due to the unilateral nature of the dynamometer configuration, only dominant limb MVCs were recorded.

However, participants also performed concurrent maximal contractions against manual resistance with the nondominant limb. The highest torque produced by participants across their four attempts was identified as their peak baseline measurement (*MVC #1*). When testing the plantar flexors, participants were seated with their lower limb near-maximally extended, their talocrural joint positioned at 0° and their foot strapped to a footplate, against which they maximally plantarflexed. When testing the knee extensors, participants were seated with their knee flexed at 90° and the distal shank strapped to a pad attached to a lever arm, against which they maximally extended. For both muscle groups, the dynamometer axis of rotation was aligned with the relevant joint center of rotation. All torque data were collected using HUMAC software (CSMi, Stoughton, MA, USA).

After *MVC #1*, participants completed an *unfatigued run* on the treadmill for 10 min at either 10 km h^-1^ [2.77 m s^-1^] (*recreational* group) or 12 km h^-1^ [3.33 m s^-1^] (*experienced* group). These speeds correspond to an easy pace and have been used when collecting steady-state metabolic data from runners of varying experience levels (66, 67). Breath-by-breath gaseous exchange was recorded by the portable metabolic system attached to a harness worn by participants (total mass: 0.9 kg). All metabolic data were collected using OMNIA software (COSMED, Rome, Italy). Electromyography data were collected continuously for all seven muscles at 2000 Hz (LabChart 8, ADInstruments, Dunedin, New Zealand). After the run, participants immediately transitioned to the dynamometer for another MVC (*MVC #2*).

Next, one of two fatigue protocols was completed depending on the session. For the plantar flexors, participants completed 2-3 sets of bilateral bodyweight calf raises on the edge of a ∼20 cm high step with 1 min rest in between sets (68). Calf raises were done to a 65-bpm metronome with a beat on full plantarflexion and a beat on full dorsiflexion. For the knee extensors, participants completed 2-3 sets of bilateral knee extensions in a knee extension machine (Hammer Strength, Rosemont, IL, USA) set to 20% of their peak torque measurement (highest torque obtained across *MVC #1* and *#2*) converted to a mass, with 1 min rest in between sets. Repetitions were completed to a 60-bpm metronome with a beat on full knee extension and a beat on full knee flexion (∼90°). Each set was done until failure, which could be volitional, or performance-based (e.g., incomplete range of motion, poor timing with the metronome). After two sets, participants immediately completed an MVC (*MVC #3*) to ensure fatigue had reached the target value of 75-85% of their peak baseline torque measure. This range of neuromuscular fatigue was chosen as it reflects the percentage torque decrease that occurs in these muscle groups after a prolonged run (6–8, 35, 43–49, 51). If fatigue was not within the suitable range, participants completed another set of calf raises or knee extensions, or rested, and fatigue was reassessed. One participant required a fourth set of calf raises but all other participants required 2-3 sets of calf raises and knee extensions. The number of sets and repetitions are found in Supplementary Tables 3 and 4. When the suitable range of fatigue was achieved, participants immediately transitioned to a *fatigued run* on the treadmill. Conditions between the *unfatigued* and *fatigued runs* were mirrored, and the same data were collected. Finally, after the *fatigued run*, participants immediately completed a final MVC (*MVC #4*). The dynamometer was in close proximity to the treadmill and fatigue protocol equipment, which allowed for a rapid measure of fatigue (∼10-20 s transition). This was important because recovery occurs rapidly after a fatiguing activity (39, 40). A summary of the protocol is presented in Figure 1.

**Figure 1.**
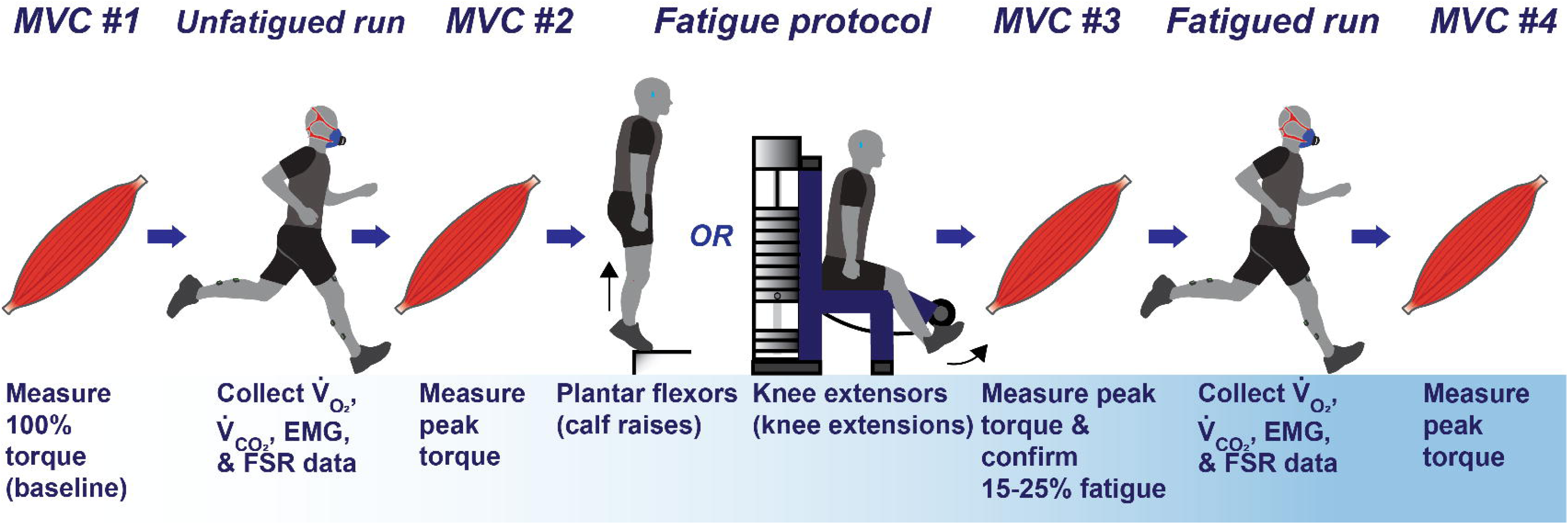
Study protocol. Conducted over two non-consecutive days, one muscle group (plantar flexors or knee extensors) per day (order randomized). MVC = maximum voluntary contraction, EMG = electromyography, FSR = force-sensing resistor.

### Data processing

All data were processed using custom Python scripts (Python Software Foundation, version 3.9). 𝑉͘_𝑂_2__ and 𝑉͘_𝐶𝑂_2__ and data were smoothed with a third-order, low pass digital filter with a cut-off frequency of 0.04 Hz (69). Resting metabolic rate was calculated from 1 min of 𝑉͘_𝑂_2__and 𝑉͘_𝐶𝑂_2__data selected from the minute with the lowest average magnitudes across the final two minutes of the seated trial, when the gaseous exchange data had reached a plateau.

To compute net metabolic power (𝐶_𝑚et,𝑃_; W kg^-1^), oxygen inspiration (𝑉͘_𝑂_ ; L s^-1^) and carbon dioxide expiration (𝑉͘_𝐶𝑂_2__; L s^-1^) resting rates were subtracted from equivalent values obtained in the *unfatigued* and *fatigued run*, and entered into Equation 1 (70, 71), where 𝑀𝑀 represents the participant’s body mass (kg):

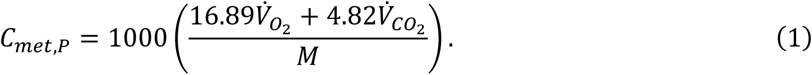

𝑉͘_𝑂_2__ and 𝑉͘_𝐶𝑂_2__ data were selected from the eighth minute of the *unfatigued* and *fatigued run* to avoid the psychological effects of participants knowing the run would end soon.

An increase in metabolic power represents worsening running economy. Even small changes (e.g., 1%) can have meaningful effects on running performance (72). However, a 4% metabolic equipment error has been observed and this must be considered when interpreting the results (73). Between-day reliability of metabolic power is unclear; however, running economy (O₂ kg⁻¹ km⁻¹) demonstrates high reliability, with an intraclass correlation coefficient of 0.81-0.83 and a typical error of 2.72-3.31% (74). Cost of transport (kcal kg⁻¹ km⁻¹) also has good-to-high reliability, with an intraclass correlation coefficient of 0.75-0.79 and a typical error of 3.05-3.72% (74). For two participants the fifth minute was used to calculate metabolic power in both runs because of supraphysiological magnitudes observed during the later stages of the run (confirmed to be a hardware error).

Respiratory exchange ratio (RER; 𝑉͘_𝐶𝑂_2__/𝑉_𝑂_2__) was calculated for the same time periods as metabolic cost. One participant’s metabolic data from both runs for both muscle groups were excluded because their RER exceeded 1.0 during the steady-state period.

Fatigue was defined as a decline in peak torque (N m) relative to the peak torque baseline measurement taken from the highest peak measurement across *MVC #1* and *#2*. Previous research has reported high between-day reliability of peak knee extensor torque measured during isometric MVC, with intraclass correlation coefficients of 0.92-0.97 and coefficients of variation of 4-5% (75). For peak plantar flexor torque measured via isometric MVC, between-subject reliability is good but shows moderate variability, with an intraclass correlation coefficient of 0.77 and a coefficient of variation value of 16.7% (76).

For the plantar flexor fatigue, 17 participants (17/18) had their baseline peak torque measurement recorded in *MVC #1* and one participant (1/18) had it recorded in *MVC #2.* For the knee extensor fatigue, 12 participants (12/18) had their baseline peak torque measurement recorded in *MVC #1* and six participants (6/18) had it recorded in *MVC #2*. For one participant, their highest knee extensor peak torque was attained during *MVC #4*. However, their *MVC #1* and *MVC #4* values fell within an appropriate intraclass correlation range for knee extension MVCs (0.92 (75)). Thus, their *MVC #1* value was selected as their baseline.

Mean and 95% confidence intervals (CI) were calculated from the predicted values derived from the linear mixed effects models (see ‘Statistical analysis’ section below). However, raw data were used when addressing atypical data points in text to illustrate their impact. Additionally, raw data were used to calculate peak torque percentage changes. The baseline measure (*MVC #1* or *#2*) was set to 100%, and subsequent MVCs were expressed as percentages relative to the baseline. Percentage changes were determined by subtracting the percentage MVC values from the baseline percentage.

Mean integrated EMG (𝑖*E𝑀G*) from *minute 1* and *minute 10* of both runs was calculated for the SOL and MG for the plantar flexor fatigue protocol and the VL and RF for the knee extensor fatigue protocol. To our understanding, there is no published data on the between-day reliability of iEMG during running when amplitudes are normalized to peak MVC signal. However, between-day reliability of plantar flexor and knee extensor root mean square EMG, normalized to the MVC signal, is moderate to high, with intraclass correlations of 0.68-0.90 and coefficients of variation of 6-8% (77).

The raw EMG data were DC offset, band-pass filtered (20-350 Hz), full-wave rectified, and low-pass filtered (6 Hz); fourth-order dual-pass Butterworth filters were used (78). The resulting linear envelopes for the *unfatigued* and *fatigued run* trials were normalized to the linear envelope of the peak EMG signal captured during *MVC #1*. The linear envelope of the peak EMG signal captured during *MVC #2* was used for normalization for two participants who did not have EMG data for *MVC #1* because of connectivity interruptions during data collection. The normalized linear envelope data (*EMG_env_*) for each muscle (𝑖 = 1-4) were then parsed into strides (𝑗 = 1, 2 …) using force-sensing resistor pad data, which were filtered with a fourth-order low-pass Butterworth filter with a 10 Hz cut-off frequency. Strides were defined from initial contact of the dominant limb (𝑡𝑡_0_) to the consecutive initial contact of the dominant limb (𝑡_𝑓_), and then converted to a rate using stride time (𝑇_𝑗_) (79, 78). The processed muscle activity data for muscle 𝑖𝑖, stride 𝑗 (𝑎_𝑖𝑗_) is represented by Equation 2:

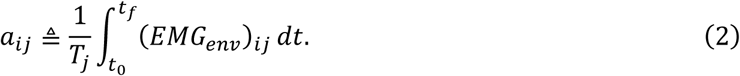

The 𝑎_𝑖𝑗_ data consisted of two one-minute windows and mean iEMG values were calculated for each window for the *unfatigued* and *fatigued run*.

Due to the time-sensitive nature of fatigue recovery, we did not stop the run or restart any part of the protocol—if an EMG sensor came loose or was moved, the affected data were excluded from the analysis. Comprehensive lists of sample sizes used to calculate study outcomes can be found in Supplementary Table 2, and lost data across the participant sample can be found in Supplementary Table 5.

### Statistical analysis

Separate linear mixed-effects models were performed to analyze data from the plantar flexor and knee extensor fatigue conditions. Participants were modeled as a random factor with a random intercept for all models. The model structures were as follows:

- Metabolic power, RER, and stride frequency: fixed effects included group (*recreational* or *experienced*), run (*unfatigued* or *fatigued*), and their interaction (two-way).
- Peak torque: fixed effects included group, MVC # (*1–4*), and their interaction (two-way).
- iEMG: fixed effects included group, run, minute (*1* and *10*), and their interactions (two- or three-way).

A likelihood ratio test was conducted to assess model fit with and without a group interaction. If model fit was improved by the inclusion of this interaction term (p<0.05), it was retained. *Pooled* group data were analyzed if the interaction was removed, while *recreational* and *experienced* group data were analyzed if the interaction was retained. Because differences between *recreational* and *experienced* runners were not relevant to the study aims, group main effects were not of interest. Differences between levels were further evaluated with post-hoc pairwise comparisons, and Bonferroni correction was applied. Custom contrasts were used for peak torque analysis (i.e., *MVC #2-1* and *MVC #4-3*) to constrain the models and investigate comparisons relevant to the study aim.

Statistical significance was set at α=0.05 and results are reported as estimated marginal mean [95% CI] and mean differences [95% CI]. R (R Development Core Team, Vienna, Austria) was used for all statistical analyses.

## RESULTS

### Metabolic power

Individual participant metabolic power data and group means for the plantar flexor and knee extensor fatigue protocols are presented in Figure 2A and Figure 2B, respectively.

**Figure 2.**
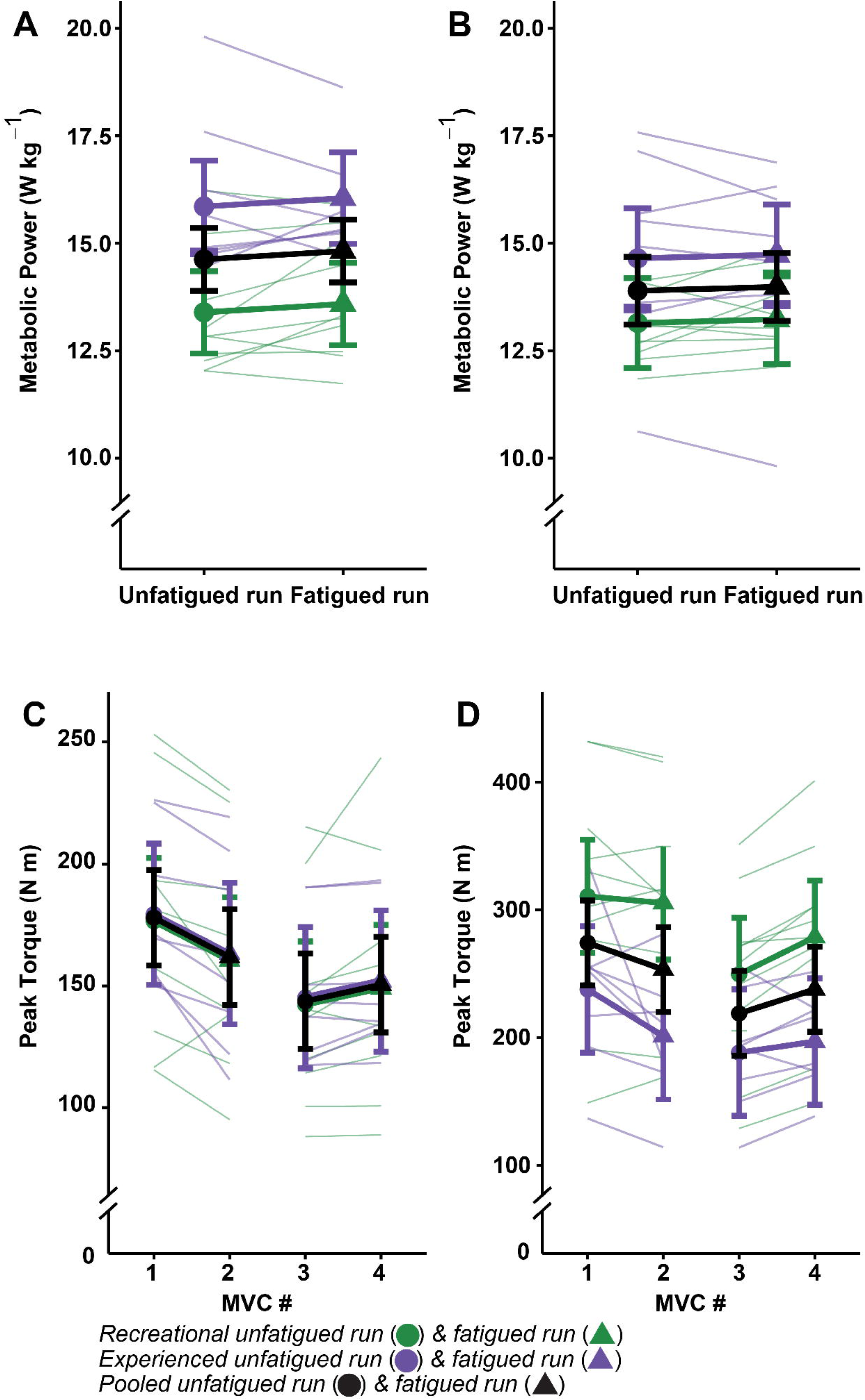
The metabolic cost of **A.** plantar flexor and **B.** knee extensor fatigue during the *unfatigued run* and *fatigued run*. **C.** Plantar flexor and **D.** knee extensor fatigue before (*MVC #1*) and after (*MVC #2*) the *unfatigued run*, as well as before (*MVC #3*) and after (*MVC #4*) the *fatigued run*. *Pooled* group (N=18) mean data in black, *recreational* group (N=10) data in green, and *experienced* group (N=8) data in purple. The mean 95% CI are represented by horizontal bolded lines. Individual participant data, colored by group affiliation, are displayed as thin lines in the background. MVC = maximum voluntary contraction.

#### Plantar flexor fatigue

The interaction between group and run did not improve model fit (p=0.125) and thus was removed. There was no change in metabolic power between the *unfatigued run* (14.6 W kg⁻¹ [13.9, 15.3]) and *fatigued run* (14.8 W kg⁻¹ [14.1, 15.5]), with a difference of 0.2 W kg⁻¹ [-0.3, 0.6] (p=0.367). Therefore, plantar flexor fatigue did not induce a change in metabolic power during running.

#### Knee extensor fatigue

The interaction between group and run did not improve model fit (p=0.072) and thus was removed. There was no change in metabolic power between the *unfatigued run* (13.9 W kg^-1^ [13.1, 14.7]) and *fatigued run* (14.0 W kg^-1^ [13.2, 14.8]), with a difference of 0.1 W kg^-1^ [-0.3, 0.4] (p=0.607). Therefore, knee extensor fatigue did not induce a change in metabolic power during running.

### Respiratory exchange ratio (RER)

#### Plantar flexor fatigue

The interaction between group and run improved model fit (p=0.023) and thus was retained. For the *recreational* group, RER was greater in the *unfatigued run* (0.88 [0.84, 0.92]) than *fatigued run* (0.86 [0.81, 0.90]), with a difference of -0.02 [-0.04, 0.00] (p=0.030). Conversely, in the *experienced* group, RER was unchanged between the *unfatigued run* (0.86 [0.82, 0.91]) and the *fatigued run* (0.87 [0.83, 0.92]), with a difference of 0.01 [-0.01, 0.03] (p=0.341). Thus, compared to unfatigued running, running with plantar flexor fatigue decreased RER in *recreational* but not *experienced* runners.

#### Knee extensor fatigue

The interaction between group and run did not improve model fit (p=0.197) and thus was removed. RER was lower in the *unfatigued run* (0.86 [0.84, 0.88]) compared to the *fatigued run* (0.84 [0.82, 0.87]), with a difference of -0.02 [-0.03, 0.00] (p=0.015). Therefore, compared to unfatigued running, running with knee extensor fatigue increased RER.

### Peak torque (fatigue)

Individual participant joint torque data and group means for the plantar flexor and knee extensor fatigue protocols are presented in Figure 2C/Table 1 and Figure 2D/Table 2, respectively.

**Table 1.** Mean peak plantar flexor torque (N m [95% CI]) for maximum voluntary contractions (MVCs) #1-4 for the pooled group (N=18), recreational group (N=10), and experienced group (N=8). Differences (N m [95% CI]) are provided for MVC #2-1 and #4-3 for the pooled group as an interaction between group and MVC # was excluded.

| Group | MVC #1 | MVC #2 | MVC #3 | MVC #4 | Difference (MVC #2-1) | p-value (MVC #2-1) | Difference (MVC #4-3) | p-value (MVC #4-3) |
| --- | --- | --- | --- | --- | --- | --- | --- | --- |
| **Pooled** | 178<br>[158, 197] | 161<br>[142, 181] | 143<br>[124, 163] | 150<br>[130, 170] | -16*<br>[-24, -8] | p<0.001* | 7*<br>[-1, 15] | p=0.033* |
| **Recreational** | 176<br>[150, 202] | 160<br>[134, 186] | 142<br>[116, 168] | 149<br>[123, 175] |  |  |  |  |
| **Experienced** | 179<br>[150, 208] | 163<br>[134, 192] | 145<br>[116, 174] | 152<br>[120, 179] |  |  |  |  |

\*Indicates statistical significance (p<0.05).

**Table 2.** Mean peak knee extensor torque (N m [95% CI]) for maximum voluntary contractions (MVCs) #1-4 for the pooled group (N=18), 815 recreational group (N=10), and experienced group (N=8). Differences (N m [95% CI]) are provided for MVC #2-1 and #4-3 for the recreational and 816 experienced group as an interaction between group and MVC # was excluded.

| Group | MVC #1 | MVC #2 | MVC #3 | MVC #4 | Difference (MVC #2-1) | p-value (MVC #2-1) | Difference (MVC #4-3) | p-value (MVC #4-3) |
| --- | --- | --- | --- | --- | --- | --- | --- | --- |
| **Pooled** | 273<br>[240, 307] | 252<br>[219, 286] | 218<br>[185, 251] | 237<br>[204, 270] |  |  |  |  |
| **Recreational** | 310<br>[266, 354] | 305<br>[260, 349] | 249<br>[204, 293] | 278<br>[234, 322] | -5<br>[-28, 18] | p=0.578 | 29*<br>[6, 52] | p=0.009* |
| **Experienced** | 237<br>[187, 286] | 200<br>[151, 250] | 187<br>[138, 237] | 196<br>[147, 246] | 37*<br>[-62, -11] | p=0.003* | 9<br>[-17, 34] | p=0.826 |

\*Indicates statistical significance (p<0.05).

#### Plantar flexor fatigue

The interaction between group and MVC # did not improve model fit (p=0.707) and thus was removed. Plantar flexor torque was greater when measured before (*MVC #1*) as compared to after (*MVC #2*) the *unfatigued run* (p<0.001). The plantar flexor fatigue protocol resulted in a 20% [19, 22] reduction in maximal torque. Plantar flexor torque increased from before (*MVC #3*) to after (*MVC #4*) the *fatigued run* (p=0.033). Therefore, plantar flexor fatigue accumulated over the *unfatigued run* and recovered over the *fatigued run*.

#### Knee extensor fatigue

The interaction between group and MVC # improved model fit (p=0.016) and thus was retained. For the *recreational* group, knee extensor torque was unchanged when measured before as compared to after the *unfatigued run* (p=0.578). The knee extensor fatigue protocol generated 21% [20, 23] fatigue in this cohort. Their knee extensor torque increased from before to after the *fatigued run* (p=0.009). For the *experienced* group, knee extensor torque was greater when measured before as compared to after the *unfatigued run* (p=0.003). The knee extensor fatigue protocol generated 22% [19, 24] fatigue in this cohort. Their knee extensor torque was unchanged when measured before and after the *fatigued run* (p=0.826). Therefore, *recreational* runners maintained stable torque production during the *unfatigued run* and recovered over the *fatigued run*, while *experienced* runners accumulated knee extensor fatigue over the *unfatigued run* and did not recover over the *fatigued run*.

### Muscle activity

Muscle activity data is presented in Figure 3.

**Figure 3.**
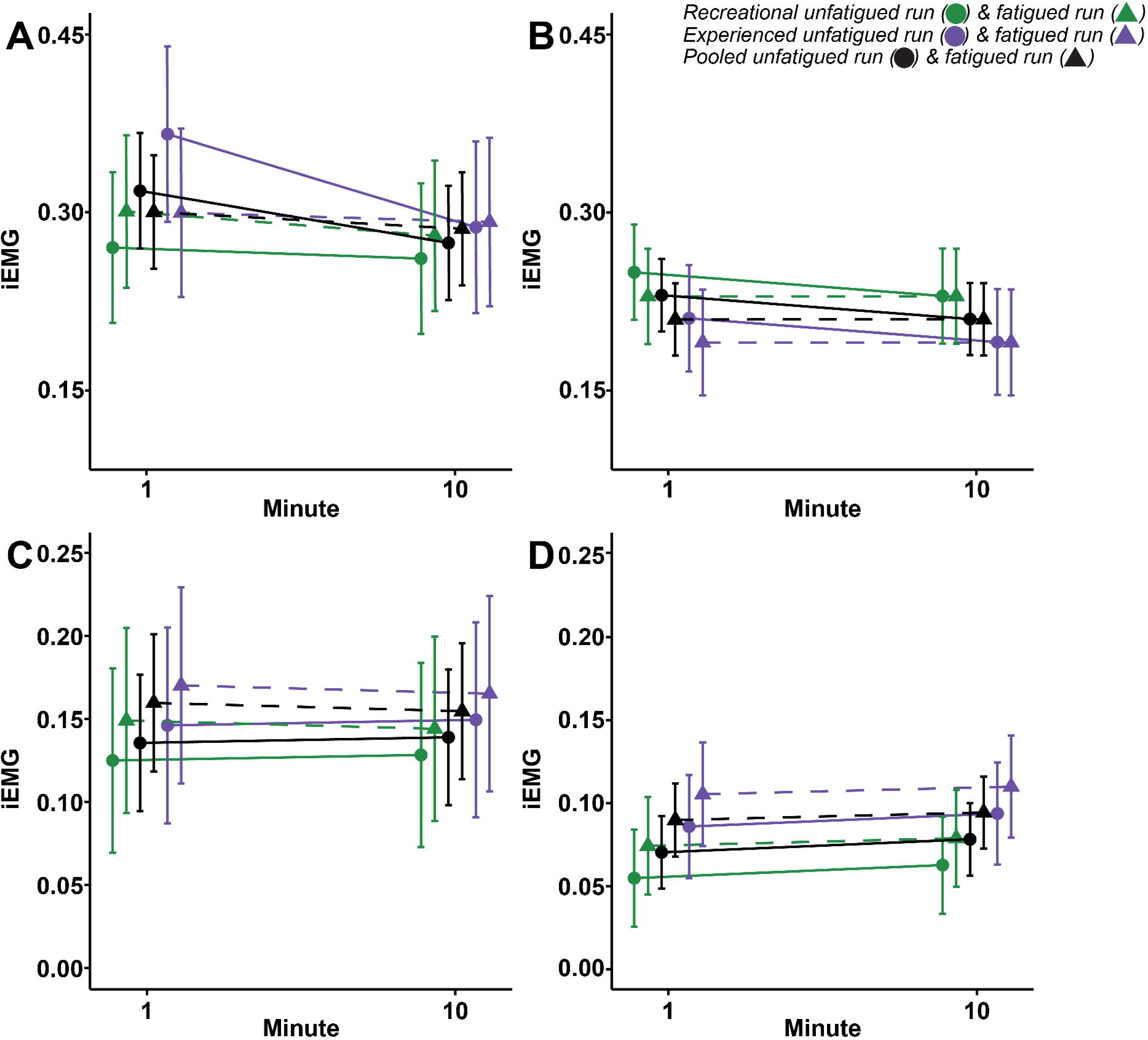
Maximum voluntary contraction (MVC)-normalized integrated mean electromyography (iEMG) measurements of the **A.** soleus (SOL) and **B.** medial gastrocnemius (MG) muscles for plantar flexor fatigue, and **C.** vastus lateralis (VL) and **D.** rectus femoris (RF) for knee extensor fatigue. The normalized linear envelope was normalized to the peak baseline MVC signal, parsed into strides (foot contact to foot contact of the same foot), presented as a rate, and averaged for *minute 1* and *minute 10* of the *unfatigued* (circles and solid lines) and *fatigued* (triangles and dashed lines) *run*. Error bars represent the 95% CI. *Pooled* (black; N=18), *recreational* (green; N=10), and *experienced* (purple; N=8) group data are presented for all muscles.

#### Plantar flexor fatigue

For SOL activity, the interactions between group and other factors improved model fit (p=0.017) and thus were retained. For the *recreational* group, SOL iEMG in *minute 1* was unchanged between the *unfatigued run* (0.27 [0.21, 0.33]) and the *fatigued run* (0.30 [0.24, 0.36]), with a difference of 0.03 [0.02, 0.08] (p=0.312). Similarly, at *minute 10*, SOL iEMG was unchanged between the *unfatigued run* (0.26 [0.20, 0.32]) and the *fatigued run* (0.28 [0.22, 0.34]), with a difference of 0.02 [-0.03, 0.07] (p=0.350). For the *experienced* group, SOL iEMG in *minute 1* was greater in the *unfatigued run* (0.37 [0.29, 0.44]) than the *fatigued run* (0.30 [0.23, 0.37]), with a difference of -0.07 [-0.13, -0.01] (p=0.022). However, a*t minute 10*, SOL iEMG was unchanged between the *unfatigued run* (0.29 [0.22, 0.36]) and the *fatigued run* (0.29 [0.22, 0.36]), with a difference of 0.00 [-0.05, 0.06] (p=0.853). Therefore, in *recreational* runners, plantar flexor fatigue did not affect SOL iEMG during running, but for the *experienced* runners, it reduced initial SOL iEMG during running.

For MG activity, the interactions between group and other factors did not improve model fit (p=0.872) and thus were removed. MG iEMG in *minute 1* of the *unfatigued run* (0.23 [0.20, 0.26]) was greater than *minute 1* of the *fatigued run* (0.21 [0.18, 0.24]), with a difference of -0.02 [-0.04, 0.00] (p=0.009).

However, at *minute 10*, MG iEMG during the *unfatigued run* (0.21 [0.18, 0.24]) was unchanged from the *fatigued run* (0.21 [0.18, 0.24]), with a difference of 0.00 [-0.02, 0.02] (p=0.969). Therefore, plantar flexor fatigue initially reduced MG iEMG during running.

#### Knee extensor fatigue

For VL activity, the interactions between group and other factors did not improve model fit (p=0.053) and thus were removed. VL iEMG in *minute 1* of the *unfatigued run* (0.14 [0.09, 0.18]) was lower than *minute 1* of the *fatigued run* (0.16 [0.12, 0.20]), with a difference of 0.02 [0.00, 0.04] (p=0.018).

However, at *minute 10*, VL iEMG during the *unfatigued run* (0.14 [0.10, 0.18]) was unchanged from the *fatigued run* (0.16 [0.11, 0.20]), with a difference of 0.02 [0.00, 0.04] (p=0.061). Therefore, knee extensor fatigue initially increased VL iEMG during running.

For RF activity, the interactions between group and other factors did not improve model fit (p=0.964) and thus were removed. RF iEMG in *minute 1* of the *unfatigued run* (0.07 [0.05, 0.09]) was lower than *minute 1* of the *fatigued run* (0.09 [0.07, 0.11]), with a difference of 0.02 [0.00, 0.03] (p=0.009).

Additionally, at *minute 10*, RF iEMG during the *unfatigued run* (0.08 [0.06, 0.10]) was lower than the *fatigued run* (0.09 [0.07, 0.12]), with a difference of 0.02 [0.00, 0.03] (p=0.010). Therefore, knee extensor fatigue increased RF iEMG during running.

### Stride frequency

#### Plantar flexor fatigue

The interaction between group and run did not improve model fit (p=0.555) and thus was removed. There was no change in stride frequency between the *unfatigued run* (82.0 strides min^-1^ [79.6, 84.4]) and *fatigued run* (82.1 strides min^-1^ [79.7, 84.6]), with a difference of 0.2 strides min^-1^ [-0.3, 0.7] (p=0.506). Therefore, running with plantar flexor fatigue did not affect stride frequency.

#### Knee extensor fatigue

The interaction between group and run did not improve model fit (p=0.531) and thus was removed. There was no change in stride frequency between the *unfatigued run* (79.8 strides min^-1^ [77.7, 81.9]) and *fatigued run* (80.4 strides min^-1^ [78.7, 82.5]), with a difference of 0.6 strides min^-1^ [-0.4, 1.6] (p=0.237). Therefore, running with knee extensor fatigue did not affect stride frequency.

## DISCUSSION

Neuromuscular fatigue, accumulated globally during a prolonged running bout, like a 10-42 km run, is thought to decrease running performance (8, 80–82) via an increase in the metabolic cost of running (5, 6, 29, 83–88). Yet, the contribution of individual muscle group fatigue to this increase is unclear. Our primary aim was therefore to quantify the effect of local i) plantar flexor and ii) knee extensor fatigue on the metabolic cost of running. Our fatigue protocol of repeated calf raises or knee extensions achieved respective torque reductions of 20% for the plantar flexor group and 21-22% for knee extensor group.

These values are similar to torque reductions induced by a prolonged run (6–8, 35, 43–51). Interestingly, local muscle fatigue of either muscle group had no effect on metabolic power during running, therefore, our first hypothesis was not supported. The majority of participants (plantar flexors: 10/18, knee extensors: 13/18) did not demonstrate a meaningful change in metabolic cost (i.e., equal to or greater than 4%), beyond that which could be explained by metabolic equipment error (73).

Our findings contrast with much of the running literature examining the effect of global muscle fatigue on the metabolic cost of prolonged running (5, 6, 29, 83–87, 89). In our study, local fatigue of the plantar flexors and knee extensors likely resulted in only minimal changes across several key variables known to influence metabolic cost, including fiber type recruitment (15–18) and substrate use (21)—as supported by the minor differences in RER (<0.03) between the *unfatigued* and *fatigued runs*, as well as metabolite concentrations and the rate of ATP production (22, 23). We also observed no changes in stride frequency with fatigue of either muscle group, even though a reduced stride frequency is often regarded as a marker of fatigue (28–30). Moreover, while a comprehensive biomechanical analysis was beyond the scope of the current study, such an analysis would likely have revealed limited gait adaptations with the addition of fatigue because adaptations away from a habitual running pattern often incur a metabolic penalty (32, 33). To explain the absence of a metabolic penalty, it is plausible that i) the cumulative burden of fatigue across multiple muscle groups is required to elicit a metabolic penalty, ii) fatigue in a single muscle group not measured in this study (e.g., dorsi flexors (90)) is required to elicit a metabolic penalty, or iii) a metabolic penalty may only be observed to occur if the local fatiguing task replicates the joint and/or musculotendon specific requirements of running.

Running relies on the stretch-shortening cycle, which involves rapid stretching and shortening of the musculotendon unit, with the muscle contracting in a quasi-isometric manor (91). In contrast, the fatigue protocols incorporated slower, isolated concentric and eccentric contractions. The stretch-shortening cycle enhances force production and reduces the metabolic cost of running, in part, by elastic energy storage and release from tendons (92–94). However, a review by (95) found little to no effect of stretch-shortening cycle fatiguing tasks (e.g., running) on Achilles tendon stiffness. Moreover, while Fletcher & MacIntosh (26) reported reduced Achilles tendon stiffness and increased metabolic cost following a 90-minute run, this was not associated with meaningful plantar flexor fatigue (3%).

Plantarflexion and knee extension fatiguing tasks can affect tendon stiffness, however these studies (24, 25) utilized static stretching (e.g., 35° of dorsiflexion for 5 min), different contraction types (e.g., isometric), and greater levels of fatigue (e.g., 36% plantar flexion fatigue, 44% knee extension fatigue) than those of the current study. Although the fatiguing protocol and stretch-shortening cycle differ, their acute effects on tendon properties appear similar. Thus, changes in tendon properties do not seem to explain the preservation of metabolic cost despite 20-22% neuromuscular fatigue.

Our muscle activity analyses of SOL and MG did not provide support for our hypothesis that iEMG of fatigued muscles would increase. With plantar flexor fatigue, we found initial reductions in SOL iEMG in the *experienced* group (∼18%), as well as initial reductions in MG iEMG in the *pooled* group (∼9%). This contrasts previous research that observed a similar magnitude of fatigue (∼20%) but no change in MG iEMG following a 10 km run (45) and an increase in MG iEMG following a marathon (5). However, the current study presents the iEMG as a rate, normalized to the peak EMG signal obtained during an MVC, whereas previous research did not, so this may have contributed to the inconsistent findings. A decrease in iEMG may suggest fatigue (96), possibly via reduced neural drive, firing rate or increased inhibitory signals. However, more likely, it represents a compensatory strategy in response to fatigue. Muscle activity is redistributed between muscles of similar function during global (31, 97–99) and local (100, 101) fatigue. Thus, plantar flexor fatigue may result in a redistribution of effort to non-fatigued muscles, such as the hip or knee extensors.

Interestingly, muscle activity analyses for VL and RF during knee extensor fatigue conflicted the plantar flexor results and did provide some support for our hypothesis that iEMG of the fatigued muscles would increase. We found initial increases in VL iEMG (∼18%), as well as initial (∼24%) and final (∼19%) increases in RF iEMG with knee extensor fatigue. This was supported by research reporting increased VL activity following a ∼2 h run at 13.80 km h^-1^ [3.82 m s^-1^] (57) and increased vastus medialis activity during running at 12.96 km h^-1^ [3.60 m s^-1^] after a simultaneous knee extensor and flexor fatigue protocol (61). Yet, VL and RF activity did not change over a five-hour run at 10.00 km h^-1^ [2.77 m s^-1^] (50). This discrepancy may stem from differences in run intensity, duration and participant fitness. The VL iEMG returned to *unfatigued run* levels in the final minute of the *fatigued run* but the RF iEMG did not.

Differences in knee extensor muscle activity may stem from differences in muscle fiber distribution or role in the gait cycle (i.e., the RF flexes the hip and extends the knee, whereas the VL only extends the knee). The muscle activity analyses may collectively indicate that plantar flexor recovery is of greater priority, consistent with their primary role in propulsion (41).

### Application to distance running

The participants in this study had ∼80% torque or strength reserves after the fatigue protocols and this may have been sufficient to meet the work demands of the fatigued run (106). Our participants’ maximum plantar flexor torque values were slightly lower, while knee extension torque values were much higher than the peak torque measures observed during similar-speed running (106). Our maximum torque was measured via isometric contractions, which do not capture the elastic energy contributions available to aid joint torque during running. The Achilles tendon plays a significant role in ankle torque generation during running (95). Therefore, despite acute plantar flexor and knee extensor fatigue (∼20%), the work demands of the run were adequately met with the strength reserves.

Considering this, runners may choose not to slow down during periods of lower limb fatigue in submaximal conditions but instead maintain the same pace knowing that their strength reserves are sufficient to maintain task demands. Nevertheless, fatigue perception does not accurately reflect neuromuscular fatigue; rather, it encompasses a combination of physiological, psychological, and sensory factors, which could make an in-run analysis of fatigue challenging.

### Considerations and limitations

The sample sizes utilized in this study are relatively low (N=20; *recreational:* 10*, experienced:* 8) therefore it is unclear whether they were sufficiently powered to detect smaller effects, and whether the effects detected were genuine or false-positive findings. Therefore, a focus on the size (estimate) and precision (95% CI) of the effects from the study could help us to better understand what results may be of interest (e.g., effects with large sizes or narrow confidence intervals), and what may be less interesting (e.g., effects with small sizes or wide confidence intervals).

For the *recreational* group, both muscle groups recovered across the *fatigued run*, which may, in part, explain the lack of metabolic penalty observed. For example, type II fibers may not have been involved during the period of metabolic cost measurement. To assess whether the use of an earlier (presumably less recovered) timepoint in the run would have impacted the results, we also analyzed metabolic cost during the fifth minute of the run; we found no effect of fatigue from either muscle on metabolic power (p<0.05). Thus, it is unlikely that metabolic power was different between the *unfatigued* and *fatigued running* conditions for either muscle group.

One participant achieved their highest maximum knee extensor torque after the fatigue run, despite being sufficiently fatigued (∼17%) before the run (Figure 2C). This may reflect a learning effect related to the MVC task, as they had no prior experience with it. In contrast, Figure 2D highlights two participants with notable knee extensor fatigue after the *fatigued run* (∼7% and ∼10% torque reductions, respectively). One participant, a habitual barefoot runner, may have lower knee extensor endurance due to a plantar flexor bias and non-rearfoot strike. The other participant showed consistent fatigue accumulation throughout the protocol, potentially influenced by unconfirmed factors like psychological stress or illness. All participants were included in the analysis, as their results were deemed valid.

## Conclusion

This study quantified the individual effects of local plantar flexor and knee extensor fatigue on the metabolic cost of running. Running with ∼20% plantar flexor and, separately, 21-22% knee extensor neuromuscular fatigue had no effect on the metabolic cost of running for *recreational* and *experienced* runners. Therefore, optimal submaximal running performance appears to be unaffected by local fatigue of these muscle groups. However, we do not discount an indirect effect of local fatigue on subjective determinants, such as rate of perceived exertion. These findings can be applied to racing and training practices, e.g., development of pacing strategies for races and strength training programs for performance.

## DATA AVAILABILITY

Source data for this study are not publicly available due to privacy or ethical restrictions. The source data are available to verified researchers upon request by contacting the corresponding author.

## ACKNOWLEDGMENTS

The authors would like to express their gratitude for the support they received from Dr Rachel Ward and Dr Martin Lindley. The authors thank Dr Ward and Dr Lindley for providing helpful feedback on earlier versions of this manuscript, also acknowledging Dr Ward’s contribution to PhD student supervision (KN), and Dr Lindley’s advice related to metabolic cost measurement. The authors also extend their thanks to Dr Guillaume Millet and Dr Kim Hérbert-Losier for their thoughtful comments on an earlier version of this work.

## DISCLAIMERS

This research was funded by ASICS Oceania^TM^; however, no ASICS Oceania^TM^ employee was directly involved in the research in any way that could have influenced or biased the reported results.

## AUTHOR CONTRIBUTIONS

KN: Conceived and designed the research, performed data collection, analyzed data, interpreted results, prepared figures, drafted the manuscript, edited and revised the manuscript, and approved the final version of the manuscript.

HF: Contributed to study design, interpreted results, revised the manuscript, and approved the final version of the manuscript.

JK: Contributed to data collection, revised the manuscript, and approved the final version of the manuscript.

MH: Contributed to study design, interpreted results, prepared figures, revised the manuscript, and approved the final version of the manuscript.

KM: Conceived and designed the research, analyzed data, interpreted results, prepared figures, revised the manuscript, and approved the final version of the manuscript.

## REFERENCES

1. Conley DL, Krahenbuhl GS. Running economy and distance running performance of highly trained athletes. Med Sci Sports Exerc 12: 357–360, 1980.

2. Joyner MJ. Modeling: optimal marathon performance on the basis of physiological factors. J Appl Physiol 70: 683–687, 1991. doi: 10.1152/jappl.1991.70.2.683.

3. Saunders PU, Pyne DB, Telford RD, Hawley JA. Factors Affecting Running Economy in Trained Distance Runners. Sports Med 34: 465–485, 2004. doi: 10.2165/00007256-200434070-00005.

4. Enoka RM, Stuart DG. Neurobiology of muscle fatigue. J Appl Physiol Bethesda Md 1985 72: 1631–1648, 1992. doi: 10.1152/jappl.1992.72.5.1631.

5. Nicol C, Komi PV, Marconnet P. Effects of marathon fatigue on running kinematics and economy. Scand J Med Sci Sports 1: 195–204, 1991. doi: 10.1111/j.1600-0838.1991.tb00296.x.

6. Petersen K, Hansen CB, Aagaard P, Madsen K. Muscle mechanical characteristics in fatigue and recovery from a marathon race in highly trained runners. Eur J Appl Physiol 101: 385–396, 2007. doi: 10.1007/s00421-007-0504-x.

7. Lloria-Varella J, Besson T, Varesco G, Espeit L, Kennouche D, Delattre N, Millet GY, Morio C, Rossi J. Running pattern changes after a 38-km trail running race: does shoe fatigue play a role? Footwear Sci 14: 185–197, 2022. doi: 10.1080/19424280.2022.2086302.

8. Ross EZ, Goodall S, Stevens A, Harris I. Time course of neuromuscular changes during running in well-trained subjects. Med Sci Sports Exerc 42: 1184–1190, 2010. doi: 10.1249/MSS.0b013e3181c91f4e.

9. Burt D, Lamb K, Nicholas C, Twist C. Effects of repeated bouts of squatting exercise on sub-maximal endurance running performance. Eur J Appl Physiol 113: 285–293, 2013. doi: 10.1007/s00421-012-2437-2.

10. Ratkevicius A, Stasiulis A, Dubininkaite L, Skurvydas A. Muscle Fatigue Increases Metabolic Costs of Ergometer Cycling without Changing VO2 Slow Component. J Sports Sci Med 5: 440–448, 2006.

11. Marcora SM, Bosio A, de Morree HM. Locomotor muscle fatigue increases cardiorespiratory responses and reduces performance during intense cycling exercise independently from metabolic stress. Am J Physiol-Regul Integr Comp Physiol 294: R874–R883, 2008. doi: 10.1152/ajpregu.00678.2007.

12. Hayes PR, French DN, Thomas K. The Effect of Muscular Endurance on Running Economy. J Strength Cond Res 25: 2464–2469, 2011. doi: 10.1519/JSC.0b013e3181fb4284.

13. Conwit RA, Stashuk D, Suzuki H, Lynch N, Schrager M, Metter EJ. Fatigue effects on motor unit activity during submaximal contractions. Arch Phys Med Rehabil 81: 1211–1216, 2000. doi: 10.1053/apmr.2000.6975.

14. Stock MS, Beck TW, Defreitas JM. Effects of fatigue on motor unit firing rate versus recruitment threshold relationships. Muscle Nerve 45: 100–109, 2012. doi: 10.1002/mus.22266.

15. Barclay CJ, Constable JK, Gibbs CL. Energetics of fast- and slow-twitch muscles of the mouse. J Physiol 472: 61–80, 1993. doi: 10.1113/jphysiol.1993.sp019937.

16. Bottinelli R, Canepari M, Reggiani C, Stienen GJ. Myofibrillar ATPase activity during isometric contraction and isomyosin composition in rat single skinned muscle fibres. J Physiol 481: 663–675, 1994. doi: 10.1113/jphysiol.1994.sp020472.

17. He Z-H, Bottinelli R, Pellegrino MA, Ferenczi MA, Reggiani C. ATP Consumption and Efficiency of Human Single Muscle Fibers with Different Myosin Isoform Composition. Biophys J 79: 945–961, 2000. doi: 10.1016/S0006-3495(00)76349-1.

18. Stienen GJ, Kiers JL, Bottinelli R, Reggiani C. Myofibrillar ATPase activity in skinned human skeletal muscle fibres: fibre type and temperature dependence. J Physiol 493: 299–307, 1996. doi: 10.1113/jphysiol.1996.sp021384.

19. Callow M, Morton A, Guppy M. Marathon fatigue: the role of plasma fatty acids, muscle glycogen and blood glucose. Eur J Appl Physiol 55: 654–661, 1986. doi: 10.1007/BF00423212.

20. Hermansen L, Hultman E, Saltin B. Muscle Glycogen during Prolonged Severe Exercise. Acta Physiol Scand 71: 129–139, 1967. doi: 10.1111/j.1748-1716.1967.tb03719.x.

21. Hargreaves M, Spriet LL. Skeletal muscle energy metabolism during exercise. Nat Metab 2: 817–828, 2020. doi: 10.1038/s42255-020-0251-4.

22. Hopkins E, Sanvictores T, Sharma S. Physiology, Acid Base Balance [Online]. In: *StatPearls*. StatPearls Publishing http://www.ncbi.nlm.nih.gov/books/NBK507807/ [13 Apr. 2024].

23. Sutton JR, Jones NL, Toews CJ. Effect of pH on Muscle Glycolysis during Exercise. Clin Sci 61: 331–338, 1981. doi: 10.1042/cs0610331.

24. Kubo K, Kanehisa H, Kawakami Y, Fukunaga T. Effects of repeated muscle contractions on the tendon structures in humans. Eur J Appl Physiol 84: 162–166, 2001. doi: 10.1007/s004210000337.

25. Kubo K, Kanehisa H, Fukunaga T. Effects of transient muscle contractions and stretching on the tendon structures in vivo. Acta Physiol Scand 175: 157–164, 2002. doi: 10.1046/j.1365-201X.2002.00976.x.

26. Fletcher JR, MacIntosh BR. Changes in Achilles tendon stiffness and energy cost following a prolonged run in trained distance runners. PLoS ONE 13: e0202026, 2018. doi: 10.1371/journal.pone.0202026.

27. Lichtwark GA, Wilson AM. Is Achilles tendon compliance optimised for maximum muscle efficiency during locomotion? J Biomech 40: 1768–1775, 2007. doi: 10.1016/j.jbiomech.2006.07.025.

28. García-Pérez JA, Pérez-Soriano P, Llana S, Martínez-Nova A, Sánchez-Zuriaga D. Effect of overground vs treadmill running on plantar pressure: Influence of fatigue. Gait Posture 38: 929–933, 2013. doi: 10.1016/j.gaitpost.2013.04.026.

29. Hunter I, Smith GA. Preferred and optimal stride frequency, stiffness and economy: changes with fatigue during a 1-h high-intensity run. Eur J Appl Physiol 100: 653–661, 2007. doi: 10.1007/s00421-007-0456-1.

30. Van Hooren B, Jukic I, Cox M, Frenken KG, Bautista I, Moore IS. The Relationship Between Running Biomechanics and Running Economy: A Systematic Review and Meta-Analysis of Observational Studies. Sports Med Auckl NZ 54: 1269–1316, 2024. doi: 10.1007/s40279-024-01997-3.

31. Hajiloo B, Anbarian M, Esmaeili H, Mirzapour M. The effects of fatigue on synergy of selected lower limb muscles during running. J Biomech 103: 109692, 2020. doi: 10.1016/j.jbiomech.2020.109692.

32. Sanno M, Willwacher S, Epro G, Brüggemann G-P. Positive Work Contribution Shifts from Distal to Proximal Joints during a Prolonged Run. Med Sci Sports Exerc 50: 2507–2517, 2018. doi: 10.1249/MSS.0000000000001707.

33. Sanno M, Epro G, Brüggemann G-P, Willwacher S. Running into Fatigue: The Effects of Footwear on Kinematics, Kinetics, and Energetics. Med Sci Sports Exerc 53: 1217–1227, 2021. doi: 10.1249/MSS.0000000000002576.

34. Place N, Millet GY. Quantification of Neuromuscular Fatigue: What Do We Do Wrong and Why? Sports Med 50: 439–447, 2020. doi: 10.1007/s40279-019-01203-9.

35. Nicol C, Komi PV, Marconnet P. Fatigue effects of marathon running on neuromuscular performance. I. Changes in muscle force and stiffness characteristics. Scand J Med Sci Sports 1: 10–17, 1991.

36. Viitasalo JHT, Komi PV. Signal characteristics of EMG during fatigue. Eur J Appl Physiol 37: 111–121, 1977. doi: 10.1007/BF00421697.

37. Dideriksen JL, Farina D, Enoka RM. Influence of fatigue on the simulated relation between the amplitude of the surface electromyogram and muscle force. Philos Trans R Soc Math Phys Eng Sci 368: 2765–2781, 2010. doi: 10.1098/rsta.2010.0094.

38. Vøllestad NK. Measurement of human muscle fatigue. J Neurosci Methods 74: 219–227, 1997. doi: 10.1016/S0165-0270(97)02251-6.

39. Carroll TJ, Taylor JL, Gandevia SC. Recovery of central and peripheral neuromuscular fatigue after exercise. J Appl Physiol 122: 1068–1076, 2017. doi: 10.1152/japplphysiol.00775.2016.

40. Froyd C, Millet GY, Noakes TD. The development of peripheral fatigue and short-term recovery during self-paced high-intensity exercise. J Physiol 591: 1339–1346, 2013. doi: 10.1113/jphysiol.2012.245316.

41. Novacheck TF. The biomechanics of running. Gait Posture 7: 77–95, 1998. doi: 10.1016/S0966-6362(97)00038-6.

42. Riddick RC, Kuo AD. Mechanical work accounts for most of the energetic cost in human running. Sci Rep 12: 645, 2022. doi: 10.1038/s41598-021-04215-6.

43. Saldanha A, Nordlund Ekblom MM, Thorstensson A. Central fatigue affects plantar flexor strength after prolonged running. Scand J Med Sci Sports 18: 383–388, 2008. doi: 10.1111/j.1600-0838.2007.00721.x.

44. Avela J, Kyröläinen H, Komi PV. Altered reflex sensitivity after repeated and prolonged passive muscle stretching. J Appl Physiol Bethesda Md 1985 86: 1283–1291, 1999. doi: 10.1152/jappl.1999.86.4.1283.

45. Finni T, Kyröläinen H, Avela J, Komi P. Maximal but not submaximal performance is reduced by constant-speed 10-km run. J Sports Med Phys Fitness 43: 411–417, 2004.

46. Nicol C, Kuitunen S, Kyröläinen H, Avela J, Komi PV. Effects of long- and short-term fatiguing stretch-shortening cycle exercises on reflex EMG and force of the tendon-muscle complex. Eur J Appl Physiol 90: 470–479, 2003. doi: 10.1007/s00421-003-0862-y.

47. Play M-C, Giandolini M, Perrin TP, Metra M, Feasson L, Rossi J, Millet GY. Soft-Tissue Vibrations and Fatigue During Prolonged Running: Does an Individualized Midsole Hardness Play a Role? Scand J Med Sci Sports 34: e14672, 2024. doi: 10.1111/sms.14672.

48. Davies CT, Thompson MW. Physiological responses to prolonged exercise in ultramarathon athletes. J Appl Physiol 61: 611–617, 1986. doi: 10.1152/jappl.1986.61.2.611.

49. Millet GY, Martin V, Lattier G, Ballay Y. Mechanisms contributing to knee extensor strength loss after prolonged running exercise. J Appl Physiol 94: 193–198, 2003. doi: 10.1152/japplphysiol.00600.2002.

50. Place N, Lepers R, Deley G, Millet GY. Time Course of Neuromuscular Alterations during a Prolonged Running Exercise. Med Sci Sports Exerc 36: 1347, 2004. doi: 10.1249/01.MSS.0000135786.22996.77.

51. Vercruyssen F, Tartaruga M, Horvais N, Brisswalter J. Effects of Footwear and Fatigue on Running Economy and Biomechanics in Trail Runners. Med Sci Sports Exerc 48: 1976–1984, 2016. doi: 10.1249/MSS.0000000000000981.

52. Hagberg JM, Mullin JP, Nagle FJ. Oxygen consumption during constant-load exercise. J Appl Physiol 45: 381–384, 1978. doi: 10.1152/jappl.1978.45.3.381.

53. Casaburi R, Wasserman K. Exercise training in pulmonary rehabilitation. N Engl J Med 314: 1509–1511, 1986. doi: 10.1056/nejm198606053142310.

54. Westerlind KC, Byrnes WC, Mazzeo RS. A comparison of the oxygen drift in downhill vs. level running. J Appl Physiol Bethesda Md 1985 72: 796–800, 1992. doi: 10.1152/jappl.1992.72.2.796.

55. Christina KA, White SC, Gilchrist LA. Effect of localized muscle fatigue on vertical ground reaction forces and ankle joint motion during running. Hum Mov Sci 20: 257–276, 2001. doi: 10.1016/s0167-9457(01)00048-3.

56. Kellis E, Liassou C. The effect of selective muscle fatigue on sagittal lower limb kinematics and muscle activity during level running. J Orthop Sports Phys Ther 39: 210–220, 2009. doi: 10.2519/jospt.2009.2859.

57. Hausswirth C, Brisswalter J, Vallier JM, Smith D, Lepers R. Evolution of electromyographic signal, running economy, and perceived exertion during different prolonged exercises. Int J Sports Med 21: 429–436, 2000. doi: 10.1055/s-2000-3832.

58. Hicks AL, Kent-Braun J, Ditor DS. Sex Differences in Human Skeletal Muscle Fatigue. Exerc Sport Sci Rev 29: 109–112, 2001.

59. Pallavi L, D Souza UJ, Shivaprakash G. Assessment of Musculoskeletal Strength and Levels of Fatigue during Different Phases of Menstrual Cycle in Young Adults. J Clin Diagn Res JCDR 11: CC11–CC13, 2017. doi: 10.7860/JCDR/2017/24316.9408.

60. Meignié A, Duclos M, Carling C, Orhant E, Provost P, Toussaint J-F, Antero J. The Effects of Menstrual Cycle Phase on Elite Athlete Performance: A Critical and Systematic Review. Front Physiol 12: 654585, 2021. doi: 10.3389/fphys.2021.654585.

61. Micklewright D, St Clair Gibson A, Gladwell V, Al Salman A. Development and Validity of the Rating-of-Fatigue Scale. Sports Med 47: 2375–2393, 2017. doi: 10.1007/s40279-017-0711-5.

62. van Melick N, Meddeler BM, Hoogeboom TJ, Nijhuis-van der Sanden MWG, van Cingel REH. How to determine leg dominance: The agreement between self-reported and observed performance in healthy adults. PLoS ONE 12: e0189876, 2017. doi: 10.1371/journal.pone.0189876.

63. Hermens HJ, Freriks B, Merletti R, Stegeman D, Blok J, Rau G, Disselhorst-Klug C, Hägg G. European Recommendations for Surface ElectroMyoGraphy.

64. **ASICS**. GEL-CUMULUS 24 [Online]. https://www.asics.com/au/en-au/gel-cumulus-24/p/AOP_1011B366-400.html [11 Mar. 2025].

65. Binboğa E, Tok S, Catikkas F, Güven Ş, Dane S. The effects of verbal encouragement and conscientiousness on maximal voluntary contraction of the triceps surae muscle in elite athletes. J Sports Sci 31: 982–988, 2013. doi: 10.1080/02640414.2012.758869.

66. de Ruiter CJ, Verdijk PWL, Werker W, Zuidema MJ, de Haan A. Stride frequency in relation to oxygen consumption in experienced and novice runners. Eur J Sport Sci 14: 251–258, 2014. doi: 10.1080/17461391.2013.783627.

67. Mendonca GV, Matos P, Correia JM. Running economy in recreational male and female runners with similar levels of cardiovascular fitness. J Appl Physiol 129: 508–515, 2020. doi: 10.1152/japplphysiol.00349.2020.

68. Hébert-Losier K, Fernandez MaR, Athens J, Kubo M, O’Neill S. A randomised crossover trial on the effects of foot starting position on calf raise test outcomes: Position does matter. The Foot 60: 102112, 2024. doi: 10.1016/j.foot.2024.102112.

69. Robergs RA, Dwyer D, Astorino T. Recommendations for Improved Data Processing from Expired Gas Analysis Indirect Calorimetry. Sports Med 40: 95–111, 2010. doi: 10.2165/11319670-000000000-00000.

70. Kipp S, Grabowski AM, Kram R. What determines the metabolic cost of human running across a wide range of velocities? J Exp Biol 221, 2018. doi: 10.1242/jeb.184218.

71. Péronnet F, Massicotte D. Table of nonprotein respiratory quotient: an update. Can J Sport Sci J Can Sci Sport 16: 23–29, 1991.

72. Kipp S, Kram R, Hoogkamer W. Extrapolating Metabolic Savings in Running: Implications for Performance Predictions. Front Physiol 10: 79, 2019. doi: 10.3389/fphys.2019.00079.

73. Perez-Suarez I, Martin-Rincon M, Gonzalez-Henriquez JJ, Fezzardi C, Perez-Regalado S, Galvan-Alvarez V, Juan-Habib JW, Morales-Alamo D, Calbet JAL. Accuracy and Precision of the COSMED K5 Portable Analyser. Front Physiol 9, 2018. doi: 10.3389/fphys.2018.01764.

74. Shaw AJ, Ingham SA, Fudge BW, Folland JP. The reliability of running economy expressed as oxygen cost and energy cost in trained distance runners. Appl Physiol Nutr Metab 38: 1268–1272, 2013. doi: 10.1139/apnm-2013-0055.

75. Zech A, Witte K, Pfeifer K. Reliability and performance-dependent variations of muscle function variables during isometric knee extension. J Electromyogr Kinesiol 18: 262–269, 2008. doi: 10.1016/j.jelekin.2006.08.013.

76. Merlet AN, Cattagni T, Cornu C, Jubeau M. Effect of knee angle on neuromuscular assessment of plantar flexor muscles: A reliability study. PLOS ONE 13: e0195220, 2018. doi: 10.1371/journal.pone.0195220.

77. Albertus-Kajee Y, Tucker R, Derman W, Lamberts RP, Lambert MI. Alternative methods of normalising EMG during running. J Electromyogr Kinesiol Off J Int Soc Electrophysiol Kinesiol 21: 579–586, 2011. doi: 10.1016/j.jelekin.2011.03.009.

78. McDonald KA, Cusumano JP, Hieronymi A, Rubenson J. Humans trade off whole-body energy cost to avoid overburdening muscles while walking. Proc Biol Sci 289: 20221189, 2022. doi: 10.1098/rspb.2022.1189.

79. Miller RH, Umberger BR, Hamill J, Caldwell GE. Evaluation of the minimum energy hypothesis and other potential optimality criteria for human running. Proc R Soc B Biol Sci 279: 1498–1505, 2012. doi: 10.1098/rspb.2011.2015.

80. Chan-Roper M, Hunter I, W Myrer J, L Eggett D, K Seeley M. Kinematic changes during a marathon for fast and slow runners. J Sports Sci Med 11: 77–82, 2012.

81. Hanley B, Smith L, Bissas A. Kinematic variations due to changes in pace during men’s and women’s 5 km road running. Int J Sports Sci Coach 6: 243–252, 2011.

82. Hill AV. The Physiological Basis of Athletic Records1. Nature 116: 544–548, 1925. doi: 10.1038/116544a0.

83. Abe D, Muraki S, Yanagawa K, Fukuoka Y, Niihata S. Changes in EMG characteristics and metabolic energy cost during 90-min prolonged running. Gait Posture 26: 607–610, 2007. doi: 10.1016/j.gaitpost.2006.12.014.

84. Glace BW, McHugh MP, Gleim GW. Effects of a 2-Hour Run on Metabolic Economy and Lower Extremity Strength in Men and Women. J Orthop Sports Phys Ther 27: 189–196, 1998. doi: 10.2519/jospt.1998.27.3.189.

85. Kyröläinen H, Pullinen T, Candau R, Avela J, Huttunen P, Komi PV. Effects of marathon running on running economy and kinematics. Eur J Appl Physiol 82: 297–304, 2000. doi: 10.1007/s004210000219.

86. Sproule J. Running economy deteriorates following 60 min of exercise at 80% VO2max. Eur J Appl Physiol 77: 366–71, 1998. doi: 10.1007/s004210050346.

87. Unhjem RJ. Changes in running economy and attainable maximal oxygen consumption in response to prolonged running: The impact of training status. Scand J Med Sci Sports 34: e14637, 2024. doi: 10.1111/sms.14637.

88. Xu F, Montgomery DL. Effect of Prolonged Exercise at 65 and 80 % of VO2max on Running Economy. Int J Sports Med 16: 309–313, 1995. doi: 10.1055/s-2007-973011.

89. Xu J, Turner A, Comyns TM, Harry JR, Chavda S, Bishop C. Countermovement Rebound Jump: A Comparison of Joint Work and Joint Contribution to the Countermovement and Drop Jump Tests. Appl Sci 13: 10680, 2023. doi: 10.3390/app131910680.

90. Ross EZ, Middleton N, Shave R, George K, Nowicky A. Corticomotor excitability contributes to neuromuscular fatigue following marathon running in man. Exp Physiol 92: 417–426, 2007. doi: 10.1113/expphysiol.2006.035972.

91. Monte A, Baltzopoulos V, Maganaris CN, Zamparo P. Gastrocnemius Medialis and Vastus Lateralis in vivo muscle-tendon behavior during running at increasing speeds. Scand J Med Sci Sports 30: 1163–1176, 2020. doi: 10.1111/sms.13662.

92. Ker RF, Bennett MB, Bibby SR, Kester RC, Alexander RM. The spring in the arch of the human foot. Nature 325: 147–149, 1987. doi: 10.1038/325147a0.

93. Monte A, Maganaris C, Baltzopoulos V, Zamparo P. The influence of Achilles tendon mechanical behaviour on “apparent” efficiency during running at different speeds. Eur J Appl Physiol 120: 2495–2505, 2020. doi: 10.1007/s00421-020-04472-9.

94. Nicol C, Avela J, Komi PV. The Stretch-Shortening Cycle. Sports Med 36: 977–999, 2006. doi: 10.2165/00007256-200636110-00004.

95. Obst SJ, Barrett RS, Newsham-West R. Immediate Effect of Exercise on Achilles Tendon Properties: Systematic Review. Med Sci Sports Exerc 45: 1534, 2013. doi: 10.1249/MSS.0b013e318289d821.

96. Komi PV, Tesch P. EMG frequency spectrum, muscle structure, and fatigue during dynamic contractions in man. Eur J Appl Physiol 42: 41–50, 1979. doi: 10.1007/BF00421103.

97. Dorel S, Drouet J-M, Couturier A, Champoux Y, Hug F. Changes of pedaling technique and muscle coordination during an exhaustive exercise. Med Sci Sports Exerc 41: 1277–1286, 2009. doi: 10.1249/MSS.0b013e31819825f8.

98. Hug F, Turpin NA, Guével A, Dorel S. Is interindividual variability of EMG patterns in trained cyclists related to different muscle synergies? J Appl Physiol Bethesda Md 1985 108: 1727–1736, 2010. doi: 10.1152/japplphysiol.01305.2009.

99. Turpin NA, Guével A, Durand S, Hug F. Fatigue-related adaptations in muscle coordination during a cyclic exercise in humans. J Exp Biol 214: 3305–3314, 2011. doi: 10.1242/jeb.057133.

100. Bouillard K, Jubeau M, Nordez A, Hug F. Effect of vastus lateralis fatigue on load sharing between quadriceps femoris muscles during isometric knee extensions. J Neurophysiol 111: 768–776, 2014. doi: 10.1152/jn.00595.2013.

101. Stutzig N, Siebert T. Muscle force compensation among synergistic muscles after fatigue of a single muscle. Hum Mov Sci 42: 273–287, 2015. doi: 10.1016/j.humov.2015.06.001.

