## supplementary data for "Submaximal running energetics are maintained despite local muscle fatigue"

### SUPPLEMENTAL MATERIAL

**Supplementary Table 1** Individual participant details. Order of muscle group (plantar flexor [PF] or knee extensor [KE]) reflects the testing order. Blank cells represent missing data.

| Group | Participant ID | Age (years) | Muscle group | Body mass (kg) | Rating of fatigue (0-10) | US men's shoe size | Height (cm) | 10 km time (mm:ss) |
|---|---|---|---|---|---|---|---|---|
| *Recreational* | S001 | 33 | PF | 69.7 | 1 | 9 | 1.80 | 48:00 |
| | | | KE | 70.1 | 1.5 | | | |
| *Recreational* | S002 | 24 | KE | 76.0 | 3 | 11 | 1.89 | 50:00 |
| | | | PF | 76.3 | 3.5 | | | |
| *Recreational* | S003 | 28 | KE | 70.6 | 6 | 10 | 1.71 | 58:00 |
| | | | PF | 70.2 | 5 | | | |
| *Recreational* | S004 | 34 | KE | 92.9 | 3 | 11 | 1.88 | 57:50 |
| | | | PF | 92.5 | 4 | | | |
| *Recreational* | S005 | 28 | PF | 89.4 | 6 | 12 | 1.84 | 58:00 |
| | | | KE | 89.3 | 4 | | | |
| *Recreational* | S006 | 40 | KE | 65.0 | 1 | 8 | 1.70 | 52:00 |
| | | | PF | 65.1 | 2 | | | |
| *Recreational* | S007 | 25 | PF | 85.0 | 4 | 12 | 1.88 | 45:00 |
| | | | KE | 85.0 | 0 | | | |
| *Recreational* | S008 | 31 | KE | 80.0 | 4 | 9 | 1.79 | 49:00 |
| | | | PF | 79.5 | 0 | | | |
| *Recreational* | S009 | 24 | PF | 80.7 | 5 | 12 | 1.91 | 41:20 |
| | | | KE | 80.0 | 2 | | | |
| *Recreational* | S010 | 28 | PF | 82.0 | 2 | 11 | 1.70 | 58:28 |
| | | | KE | 82.1 | 2 | | | |
| *Recreational* | S011 | 18 | PF | 72.0 | 3 | 12 | 1.82 | 43:00 |
| | | | KE | 73.3 | 2 | | | |
| *Experienced* | S001 | 40 | PF | 64.9 | 2 | 9 | 1.75 | 38:40 |
| | | | KE | 64.5 | 3 | | | |
| *Experienced* | S002 | 31 | KE | 81.8 | 2 | 11 | 1.83 | 38:00 |
| | | | PF | 81.0 | 1 | | | |
| *Experienced* | S003 | 20 | PF | 71.8 | 3 | 10 | 1.87 | 35:00 |
| | | | KE | 71.5 | 2 | | | |
| *Experienced* | S004 | 25 | PF | 67.5 | 2 | 9 | 1.78 | 37:00 |
| | | | KE | 67.5 | 2 | | | |
| *Experience* | S005 | 29 | KE | 77.5 | 2 | 10 | 1.78 | 35:00 |
| | | | PF | 77.5 | 2 | | | |
| *Experienced* | S006 | 19 | KE | 61.5 | 4 | 9 | 1.75 | 39:00 |
| | | | PF | 60.8 | 4 | | | |
| *Experienced* | S007 | 28 | PF | 77.7 | 0 | 11 | 1.73 | 37:00 |
| | | | KE | 76.5 | 0.5 | | | |
| *Experienced* | S008 | 23 | KE | 60.2 | | 9 | 1.78 | 35:00 |
| | | | PF | 59.8 | 7 | | | |
| *Experienced* | S009 | 33 | PF | 75.0 | | 11 | 1.84 | 37:00 |

**Supplementary Table 2** Sample sizes used to calculate metabolic power, respiratory exchange ratio (RER), fatigue, integrated electromyography (iEMG) for the plantar flexor (PF; soleus and medial gastrocnemius) and knee extensor (KE; vastus lateralis and rectus femoris) muscles, and stride frequency (SF) for the pooled, recreational and experienced groups for the PF and KE sessions.

| Muscle group | Group | Metabolic power | RER | Fatigue | iEMG | SF |
|---|---|---|---|---|---|---|
| PF | *Pooled* | 18 | 18 | 18 | 18 | 18 |
| | *Recreational* | 10 | 10 | 10 | 10 | 10 |
| | *Experienced* | 8 | 8 | 8 | 8 | 8 |
| KE | *Pooled* | 18 | 18 | 18 | 17 | 17 |
| | *Recreational* | 10 | 10 | 10 | 9 | 9 |
| | *Experienced* | 8 | 8 | 8 | 8 | 8 |

**Supplementary Table 3** Plantar flexor fatigue protocol sets and total repetitions. The protocol consisted of bilateral bodyweight calf raises on the edge of a ~20 cm high step with 1 min rest in between sets. Calf raises were done to a 65-bpm metronome with a beat on full plantarflexion and a beat on full dorsiflexion

| Group | Participant ID | Set 1 | Set 2 | Set 3 | Set 4 | Total reps |
|---|---|---|---|---|---|---|
| Recreational | S001 | 149 | 70 | | | 219 |
| Recreational | S002 | 69 | 63 | 60 | | 192 |
| Recreational | S003 | 106 | 82 | 40 | | 228 |
| Recreational | S004 | 59 | 41 | 39 | | 139 |
| Recreational | S005 | 61 | 106 | 68 | 60 | 295 |
| Recreational | S006 | 67 | 69 | | | 136 |
| Recreational | S007 | 88 | 100 | 61 | | 249 |
| Recreational | S008 | 50 | 47 | | | 97 |
| Recreational | S009 | 77 | 45 | | | 122 |
| Recreational | S010 | 65 | 70 | 34 | | 169 |
| Recreational | S011 | 110 | 72 | 48 | | 230 |
| Experienced | S001 | 90 | 60 | 33 | | 183 |
| Experienced | S002 | 30 | 31 | 24 | | 85 |
| Experienced | S003 | 86 | 89 | 70 | | 245 |
| Experienced | S004 | 65 | 49 | 31 | | 145 |
| Experienced | S005 | 60 | 52 | 22 | | 134 |
| Experienced | S006 | 61 | 50 | | | 111 |
| Experienced | S007 | 70 | 60 | 55 | | 185 |
| Experienced | S008 | 97 | 81 | 79 | | 257 |
| Experienced | S009 | 111 | 98 | | | 209 |

**Supplementary Table 4** Knee extensor fatigue protocol; mass, sets and repetitions. The protocol consisted of bilateral knee extensions in a knee extension machine set to 20% of their peak torque measurement (highest torque obtained across MVC #1 and #2) converted to a mass, with 1 min rest in between sets. Knee extensions were done to a 60-bpm metronome with a beat on full extension and a beat on flexion.

| Group | Participant ID | Mass (kg) | Set 1 | Set 2 | Set 3 | Total reps |
|---|---|---|---|---|---|---|
| Recreational | S001 | 37.5 | 45 | 50 | | 95 |
| Recreational | S002 | 41.0 | 26 | 22 | 18 | 66 |
| Recreational | S003 | 26.0 | 28 | 31 | | 59 |
| Recreational | S004 | 50.0 | 22 | 20 | | 42 |
| Recreational | S005 | 49.5 | 30 | 20 | 10 | 60 |
| Recreational | S006 | 24.0 | 23 | 22 | 21 | 66 |
| Recreational | S007 | 47.0 | 31 | 23 | 16 | 70 |
| Recreational | S008 | 35.0 | 36 | 28 | 23 | 87 |
| Recreational | S009 | 36.0 | 27 | 47 | | 74 |
| Recreational | S010 | 24.0 | 20 | 28 | | 48 |
| Recreational | S011 | 31.0 | 30 | 26 | | 56 |
| Experienced | S001 | 19.0 | 29 | 33 | | 62 |
| Experienced | S002 | 38.0 | 34 | 36 | 31 | 101 |
| Experienced | S003 | 33.0 | 27 | 25 | 20 | 72 |
| Experienced | S004 | 26.0 | 36 | 30 | 25 | 91 |
| Experienced | S005 | 40.0 | 33 | 26 | 19 | 78 |
| Experienced | S006 | 37.0 | 32 | 30 | | 62 |
| Experienced | S007 | 26.0 | 30 | 35 | | 65 |
| Experienced | S008 | 26.0 | 47 | 45 | | 92 |

**Supplementary Table 5** Lost or excluded metabolic power, respiratory exchange ratio (RER), integrated electromyography (iEMG) of the soleus (SOL), medial gastrocnemius (MG), vastus lateralis (VL), and rectus femoris (RF), as well as stride frequency (SF) data for the recreational and experienced groups over the unfatigued (UFR) and fatigued run (FR) for the plantar flexor (PF) and knee extensor (KE) sessions. Additionally, lost or excluded PF and KE torque data over maximum voluntary contraction (MVC) #1-4 for the recreational and experienced groups for the PF and KE sessions. OMNIA (COSMED, Rome, Italy) refers to the metabolic cart software. Hardware refers to the portable metabolic cart (K5, COSMED, Rome, Italy). Sensor = electromyography sensor. FSR = force-sensing resistor.

| Group | Participant | Muscle group | Run/MVC # | Variable | Reason |
|---|---|---|---|---|---|
| Recreational | S001 | KE | FR | 1 min VL iEMG | FSR sensor error |
| Recreational | S001 | KE | FR | 1 min RF iEMG | FSR sensor error |
| Recreational | S002 | PF | FR | 1 min SOL iEMG | FSR sensor error |
| Recreational | S002 | PF | FR | 1 min MG iEMG | FSR sensor error |
| Recreational | S004 | PF | MVC #2 | Peak torque | Dynamometer error |
| Recreational | S005 | KE | UFR | 1 min RF iEMG | EMG sensor error |
| Recreational | S005 | KE | UFR | 10 min RF iEMG | EMG sensor error |
| Recreational | S006 | KE | UFR | SF | FSR sensor error |

| | | | | | |
|---|---|---|---|---|---|
| Recreational | S006 | KE | FR | SF | FSR sensor error |
| Recreational | S006 | KE | UFR | 1 min VL iEMG | FSR sensor error |
| Recreational | S006 | KE | UFR | 1 min RF iEMG | FSR sensor error |
| Recreational | S006 | KE | UFR | 10 min VL iEMG | FSR sensor error |
| Recreational | S006 | KE | UFR | 10 min RF iEMG | FSR sensor error |
| Recreational | S006 | KE | FR | 1 min VL iEMG | FSR sensor error |
| Recreational | S006 | KE | FR | 1 min RF iEMG | FSR sensor error |
| Recreational | S006 | KE | FR | 10 min VL iEMG | FSR sensor error |
| Recreational | S006 | KE | FR | 10 min RF iEMG | FSR sensor error |
| Recreational | S010 | PF | UFR | Metabolic power/RER | RER > 1 |
| Recreational | S010 | PF | FR | Metabolic power/RER | RER > 1 |
| Recreational | S010 | KE | UFR | Metabolic power/RER | RER > 1 |
| Recreational | S010 | KE | FR | Metabolic power/RER | RER > 1 |
| Recreational | S010 | PF | MVC #1-4 | Peak torque | RER > 1 |
| Recreational | S010 | PF | MVC #1-4 | Peak torque | RER > 1 |
| Recreational | S010 | KE | MVC #1-4 | Peak torque | RER > 1 |
| Recreational | S010 | KE | MVC #1-4 | Peak torque | RER > 1 |
| Recreational | S010 | PF | UFR | 1 min SOL iEMG | RER > 1 |
| Recreational | S010 | PF | UFR | 1 min MG iEMG | RER > 1 |
| Recreational | S010 | PF | UFR | 10 min SOL iEMG | RER > 1 |
| Recreational | S010 | PF | UFR | 10 min MG iEMG | RER > 1 |
| Recreational | S010 | PF | FR | 1 min SOL iEMG | RER > 1 |
| Recreational | S010 | PF | FR | 1 min MG iEMG | RER > 1 |
| Recreational | S010 | PF | FR | 10 min SOL iEMG | RER > 1 |
| Recreational | S010 | PF | FR | 10 min MG iEMG | RER > 1 |
| Recreational | S010 | KE | UFR | 1 min VL iEMG | RER > 1 |
| Recreational | S010 | KE | UFR | 1 min RF iEMG | RER > 1 |
| Recreational | S010 | KE | UFR | 10 min VL iEMG | RER > 1 |
| Recreational | S010 | KE | UFR | 10 min RF iEMG | RER > 1 |
| Recreational | S010 | KE | FR | 1 min VL iEMG | RER > 1 |
| Recreational | S010 | KE | FR | 1 min RF iEMG | RER > 1 |
| Recreational | S010 | KE | FR | 10 min VL iEMG | RER > 1 |
| Recreational | S010 | KE | FR | 10 min RF iEMG | RER > 1 |
| Recreational | S010 | PF | UFR | SF | RER > 1 |
| Recreational | S010 | PF | FR | SF | RER > 1 |
| Recreational | S010 | KE | UFR | SF | RER > 1 |
| Recreational | S010 | KE | FR | SF | RER > 1 |
| Experienced | S001 | PF | UFR | Metabolic power/RER | Metabolic software error |
| Experienced | S001 | PF | FR | Metabolic power/RER | Metabolic software error |
| Experienced | S001 | PF | MVC #1-4 | Peak torque | Metabolic software error |
| Experienced | S001 | PF | MVC #1-4 | Peak torque | Metabolic software error |
| Experienced | S001 | PF | UFR | 1 min SOL iEMG | Metabolic software error |
| Experienced | S001 | PF | UFR | 1 min MG iEMG | Metabolic software error |
| Experienced | S001 | PF | UFR | 10 min SOL iEMG | Metabolic software error |

| | | | | | |
|---|---|---|---|---|---|
| Experienced | S001 | PF | UFR | 10 min MG iEMG | Metabolic software error |
| Experienced | S001 | PF | FR | 1 min SOL iEMG | Metabolic software error |
| Experienced | S001 | PF | FR | 1 min MG iEMG | Metabolic software error |
| Experienced | S001 | PF | FR | 10 min SOL iEMG | Metabolic software error |
| Experienced | S001 | PF | FR | 10 min MG iEMG | Metabolic software error |
| Experienced | S001 | PF | UFR | SF | Metabolic software error |
| Experienced | S001 | PF | FR | SF | Metabolic software error |
| Experienced | S002 | PF | UFR | 1 min SOL iEMG | EMG sensor error |
| Experienced | S002 | PF | UFR | 1 min MG iEMG | EMG sensor error |
| Experienced | S002 | KE | FR | 1 min VL iEMG | EMG sensor error |
| Experienced | S002 | KE | FR | 1 min RF iEMG | EMG sensor error |
| Experienced | S006 | PF | UFR | Metabolic power/RER* | Metabolic hardware error |
| Experienced | S006 | PF | FR | Metabolic power/RER* | Metabolic hardware error |
| Experienced | S006 | KE | UFR | 1 min VL iEMG | FSR sensor error |
| Experienced | S006 | KE | UFR | 1 min RF iEMG | FSR sensor error |
| Experienced | S007 | KE | UFR | Metabolic power/RER* | Metabolic hardware error |
| Experienced | S007 | KE | FR | Metabolic power/RER* | Metabolic hardware error |
| Experienced | S009 | PF | UFR | 1 min SOL iEMG | EMG sensor error |
| Experienced | S009 | PF | UFR | 10 min SOL iEMG | EMG sensor error |
| Experienced | S009 | KE | UFR | Metabolic power/RER | Missed session |
| Experienced | S009 | KE | FR | Metabolic power/RER | Missed session |
| Experienced | S009 | KE | MVC #1-4 | Peak torque | Missed session |
| Experienced | S009 | KE | UFR | 1 min VL iEMG | Missed session |
| Experienced | S009 | KE | UFR | 1 min RF iEMG | Missed session |
| Experienced | S009 | KE | UFR | 10 min VL iEMG | Missed session |
| Experienced | S009 | KE | UFR | 10 min RF iEMG | Missed session |
| Experienced | S009 | KE | FR | 1 min VL iEMG | Missed session |
| Experienced | S009 | KE | FR | 1 min RF iEMG | Missed session |
| Experienced | S009 | KE | FR | 10 min VL iEMG | Missed session |
| Experienced | S009 | KE | FR | 10 min RF iEMG | Missed session |
| Experienced | S009 | KE | UFR | SF | Missed session |
| Experienced | S009 | KE | FR | SF | Missed session |

*The fifth minute of metabolic data was used to calculate metabolic power in both runs because of supraphysiological magnitudes observed during the later stages of the run (confirmed to be a hardware error).
